# Cortical Depth-Dependent Modeling of Visual Hemodynamic Responses

**DOI:** 10.1101/2020.03.16.993154

**Authors:** T.C. Lacy, P.A. Robinson, K.M. Aquino, J.C. Pang

## Abstract

A physiologically based three-dimensional (3D) hemodynamic model is used to predict the experimentally observed blood oxygen level dependent (BOLD) responses versus the cortical depth induced by visual stimuli. Prior 2D approximations are relaxed in order to analyze 3D blood flow dynamics as a function of cortical depth. Comparison of the predictions with experimental data for typical stimuli demonstrates that the full 3D model matches at least as well as previous approaches while requiring significantly fewer assumptions and model parameters (e.g., there is no more need to define depth-specific parameter values for spatial spreading, peak amplitude, and hemodynamic velocity).

## 1. Introduction

Functional MRI (fMRI) is widely used to indirectly capture brain function through changes in blood flow and oxygenation that accompany changes in neural activity, via the blood oxygen level dependent (BOLD) signal. The biophysical and physiological mechanisms underlying the BOLD signal have been studied previously (Buxton et al., 1998; Friston et al., 2000; Kim and Ogawa, 2012), which have been incorporated in a cortical spatiotemporal hemodynamic model that can accurately predict induced BOLD responses to a stimulus (Drysdale et al., 2010; Aquino et al., 2012, 2014). The model has shown that certain stimuli can induce both local responses and hemodynamic waves that propagate throughout the cortex due to the intrinsic spatial couplings present between adjacent regions (Aquino et al., 2012; Lacy et al., 2016; Pang et al., 2016, 2018).

In early analyses of the abovementioned model, several approximations were made in order to simplify the calculations. One of the most important of these approximations was to simplify the three-dimensional (3D) structure of the cortex into a 2D cortical sheet by averaging the BOLD response over the cortical depth (Aquino et al., 2012, 2014). At that time, fMRI resolutions were not fine enough to resolve cortical depths reliably, so averaging over the different depths not only made the resulting calculations much more tractable but also did not affect the accuracy of the predictions of the model compared to experimental data. Moreover, it was able to produce key findings such as astrocyte-induced hemodynamic time delays (Pang et al., 2017), origins of resting-state fMRI spectrum (Pang and Robinson, 2019), techniques for imaging ocular dominance and orientation preference maps (De Oliveira et al., 2019), and improvements in population receptive field estimation (Infanti and Schwarzkopf, 2020).

However, recent advances in MRI technology, including increased accessibility to ultra-high field human scanners (7T and above), have improved experimental designs to achieve submillimeter voxel resolution (Balchandani and Naidich, 2015). Hence, it is now possible to measure the BOLD response as a function of cortical depth *z* called laminar fMRI [e.g., (Koopmans et al., 2010; De Martino et al., 2013; Kashyap et al., 2018; Finn et al., 2019; Huber et al., 2020)], but challenges remain regarding accurate interpretations of the underlying neurophysiological mechanisms of the obtained BOLD responses. This can be addressed by developing a laminar-specific hemodynamic model. Recent studies have attempted to develop this type of hemodynamic model. For example, Markuerkiaga et al. (2016) proposed a two-compartment BOLD model on top of a vascular model of the cortex to predict the laminar specificity of the BOLD signal. On the other hand, Heinzle et al. (2016) proposed a balloon-based model with two cortical depths and Havlicek and Uludag? (2020) proposed a multi-compartment balloon-based model. These two models predict temporal variations in the BOLD signal across the cortical depth, disregarding any spatial effects. Hence, they lack the ability to explain hemodynamic waves in the brain (Aquino et al., 2012; Gao et al., 2015; Gravel et al., 2017; Hindriks et al., 2019). This is addressed by Puckett et al. (2016) who proposed distinct spatiotemporal hemodynamic response function per cortical depth. Their model was based on our cortical hemodynamic model but introduced advancements by not implementing averaging over *z*, allowing estimations of BOLD versus levels of the cortical depth and with each level defined by independent spatiotemporal hemodynamic processes. This has natural advantages especially for determining when the BOLD response peaks in certain layers because the main arterial inflows of blood are localized close to the surface (Lauwers et al., 2008). For example, because the full thickness of the cortex is approximately 3 mm, and the speed of hemodynamic waves is in the order of 1–4 mm/s (Aquino et al., 2014), this means that there should be a noticeable delay in the peak in the BOLD response for layers other than layer IV. In addition, modeling the full cortical thickness allows for the properties of the cortex to vary between layers, instead of assuming that it is homogeneous, as is done when averaging over *z*.

In this paper, we extend the work of Puckett et al. (2016) by explicitly modeling the BOLD response versus the cortical depth with each *z* level interacting in space and time as a continuum. This allows for better physiological understanding of the depth-dependence of the response, and for a better determination of how the physical properties of the brain (such as the elasticity of the blood vessels) vary with *z*. We also demonstrate the accuracy of the predictions by comparing the predicted response with experimental data and fits from Puckett et al. (2016).

## 2. Theory and Methods

In this section, the physical principles used to derive the hemodynamic model linking visual stimuli with the measured BOLD response are stated, along with the equations derived from them, and the new generalizations to model the response throughout the cortical layers are explained.

### 2.1. Hemodynamic Model

Here we outline the model used, whose detailed derivation and discussion have been presented elsewhere (Drysdale et al., 2010; Aquino et al., 2012; Lacy et al., 2016; Pang et al., 2018), and explain briefly how each equation is derived from physiology.

Neural activity drives hemodynamic activity. Here, we assume a profile for the neural drive and then determine the BOLD response *Y*(**R**, *t*) at position **R** and time *t*, which can be measured in an fMRI scanner if a stimulus that produces that drive is applied. This can then be compared to experimentally induced drives.

Physically, we consider a block of cortical tissue shown in Fig. 1. The main blood inflows are assumed to occur at *≈*0.8 mm below the surface (Lauwers et al., 2008). The cortical tissue is approximated as a poroelastic medium perfused by blood (Drysdale et al., 2010). This means that the local concentration of blood within the cortex is related to the local pressure (Wang, 2000). In addition, small-scale effects (smaller than approximately 0.5 mm in linear scale) are averaged over, which thus makes our model a mean-field model. Hence, our model focuses on the mean properties of the cortical tissue and does not separately distinguish dynamics within individual vessels (e.g., arterioles and venules) or the intricacies of the vasculature. This allows us to explore generic principles that can lead to net hemodynamic effects at the scales of current fMRI voxels.

**Figure 1:**
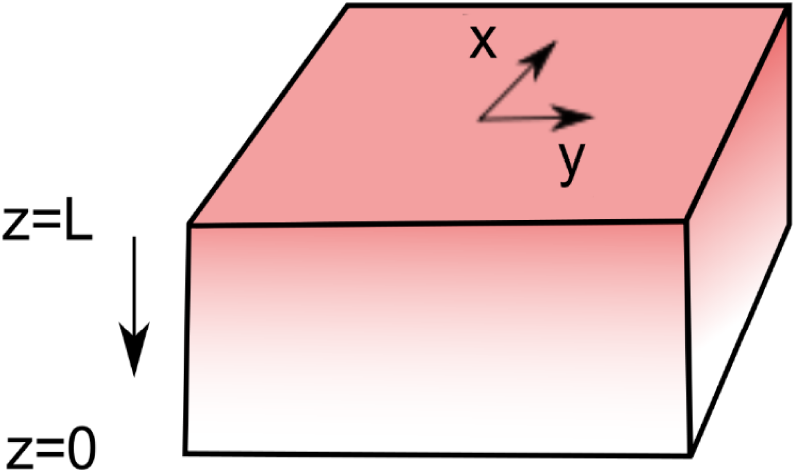
Cross-section of the block of cortical tissue, showing how the coordinates are defined, with *x* and *y* in the direction of the cortical surface and *z* perpendicular to it. The cortical surface is defined to be at *z* = *L*, while the gray-white matter boundary is at *z* = 0.

The arterial blood inflow rate (Buxton, 2009) *F* (**R**, *t*) is modeled as having a damped harmonic response driven by the neural drive *ζ*(**R**, *t*), with (Friston et al., 2000; Drysdale et al., 2010; Ress et al., 2009)

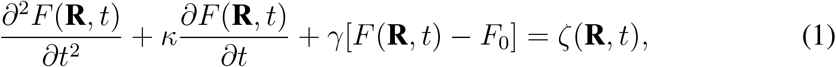

where *κ, ω*_*f*_, and *F*_0_ are the blood flow signal decay rate, the natural frequency of flow response, and the arterial blood inflow with no applied stimulus, respectively.

The total mass density of the brain is the sum of contributions due to the brain tissue and blood. The total blood mass per tissue volume Ξ(**R**,*t*) is calculated by relating it to the blood pressure using conservation of mass and momentum of blood within the tissue. The pressure within blood vessels *P* (**R**, *t*) is related to Ξ(**R**,*t*) via the constitutive equation (Aquino et al., 2012)

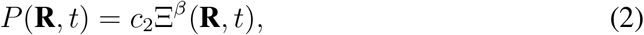

where *β* is the elasticity exponent of the blood vessels and *c*_2_ is an empirically derived constant of proportionality. This gives the pressure in terms of blood mass density, assuming that the tissue is a continuum. Note that the pressure here is the mean local pore pressure in the vasculature. Hence, it does not separately quantify pressures within individual vascular units (e.g., descending arteries) and their effects on neighboring vascular networks. A value of *β* = 1 corresponds to the vessels being elastic, while a value of *β ≈* 3.23 used here [from the inverse of Grubb’s exponent, which relates cerebral blood flow to cerebral blood volume (Grubb et al., 1974)] implies that the vessels are hyperelastic (i.e., more resistant to stretching than perfectly elastic vessels). The value of *β* depends on the type of vessel being considered (arterioles are more elastic than veins, and so have a higher *β*). Hence, since our model is a meanfield model, we can take an average value for all vessels.

Conservation of blood mass within the brain implies (Aquino et al., 2014)

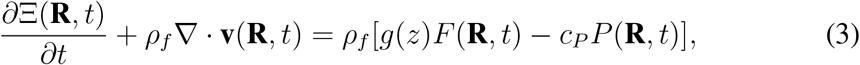

which relates the rate of change of the blood mass within the cortical tissue to inflows *ρ*_*f*_ [*g*(*z*)*F* (**R**, *t*)] and outflows *-ρ*_*f*_ [*c*_*P*_ *P* (**R**, *t*)] of blood. *ρ*_*f*_ is the blood mass density and *c*_*P*_ is the blood outflow constant. **v**(**R**, *t*) is the average velocity of blood across the vasculature. The inflow function *g*(*z*) is chosen to be a Gaussian centered at *z* = 2.4 mm to reflect the higher level of inflow near that depth with full width at half maximum (FWHM) of approximately 1 mm (Blasdel and Campbell, 2001). A Gaussian inflow function also follows Puckett et al. (2016) and is a fair first approximation since we do not have precise knowledge of the underlying neuronal inputs vs. cortical depth. The dependence of Ξ(**R**, *t*) at equilibrium on *z* is determined by measuring the total vascular volume (Lauwers et al., 2008) and then finding a profile that matches the measured values.

Momentum is also conserved for the blood. By equating the forces acting on the blood with the rate of change of momentum density, one obtains

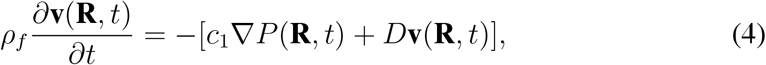

where *c*_1_ is the pressure coupling constant and *D* is the effective blood viscosity. Note that blood viscosity could potentially also change as a function of vasodilation; however, we omit this second-order effect in the present work.

From Eqs (3) and (4), one obtains the following nonlinear equation for Ξ(**R**,*t*)(Aquino et al., 2012, 2014):

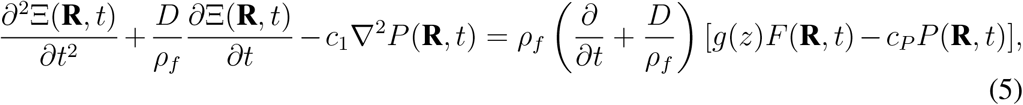

which relates Ξ(**R**,*t*) to *F*(**R**,*t*) because *P*(**R**,*t*) is a function of Ξ(**R**,*t*) via Eq. (2).

The concentration of deoxyhemoglobin (dHb) *Q*(**R**,*t*) can be determined by relating its time derivative to its flux into nearby regions, its rate of increase from the consumption of oxygen from oxyhemoglobin (oHb) by neurons, and its rate of decrease due to outflows of blood (Drysdale et al., 2010). This gives an equation that relates *Q*(**R**,*t*) to Ξ(**R**,*t*) and the boundary conditions of the system, with the rate of flow out of the cortical tissue determining how fast dHb is cleared from the brain. So, hemoglobin conservation yields (Lacy et al., 2016)

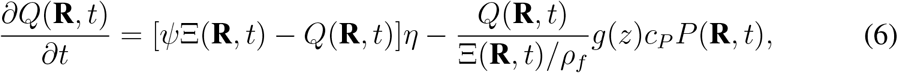

where *Ψ* is the ratio of hemoglobin concentration to blood density in mol/kg in SI units and *η* is the fractional oxygen consumption rate per unit time in s^*-*1^ in SI units. As in Lacy et al. (2016), we assume that the divergence term, which is needed in a typical continuity/conservation equation, can be neglected because the blood flows within the tissue will mostly be from arterioles and will contain very little dHb. This assumption is based on works showing that vein diameters do not change considerably relative to arteriole diameters during stimulations of less than *≈*30 s (Vazquez et al., 2010; Drew et al., 2011); hence, changes in blood mass density are primarily confined to arterioles.

Finally, the BOLD signal *Y*(**R**,*t*) is determined from Ξ(**R**,*t*) and *Q*(**R**,*t*) using a semiempirical relation derived from the properties of the fMRI scanner such as the magnetic field strength and the method of signal acquisition (Stephan et al., 2007). This gives

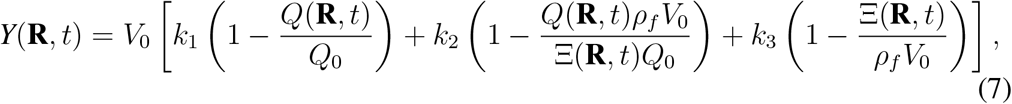

where *V*_0_ is the resting blood volume fraction, *Q*_0_ is the resting dHb concentration per unit volume, and *k*_1_, *k*_2_, and *k*_3_ are parameters that depend on the fMRI scanner being used and the experimental protocol (Obata et al., 2004; Stephan et al., 2007). Note that the nominal values of all model parameters were found elsewhere (Aquino et al., 2012, 2014; Pang et al., 2017, 2018) and are summarized in Table 1. Moreover, the readers are advised to refer to our previous articles for detailed discussions and derivations of the model equations (Drysdale et al., 2010; Aquino et al., 2012, 2014; Pang et al., 2016; Lacy et al., 2016).

**Table 1:**
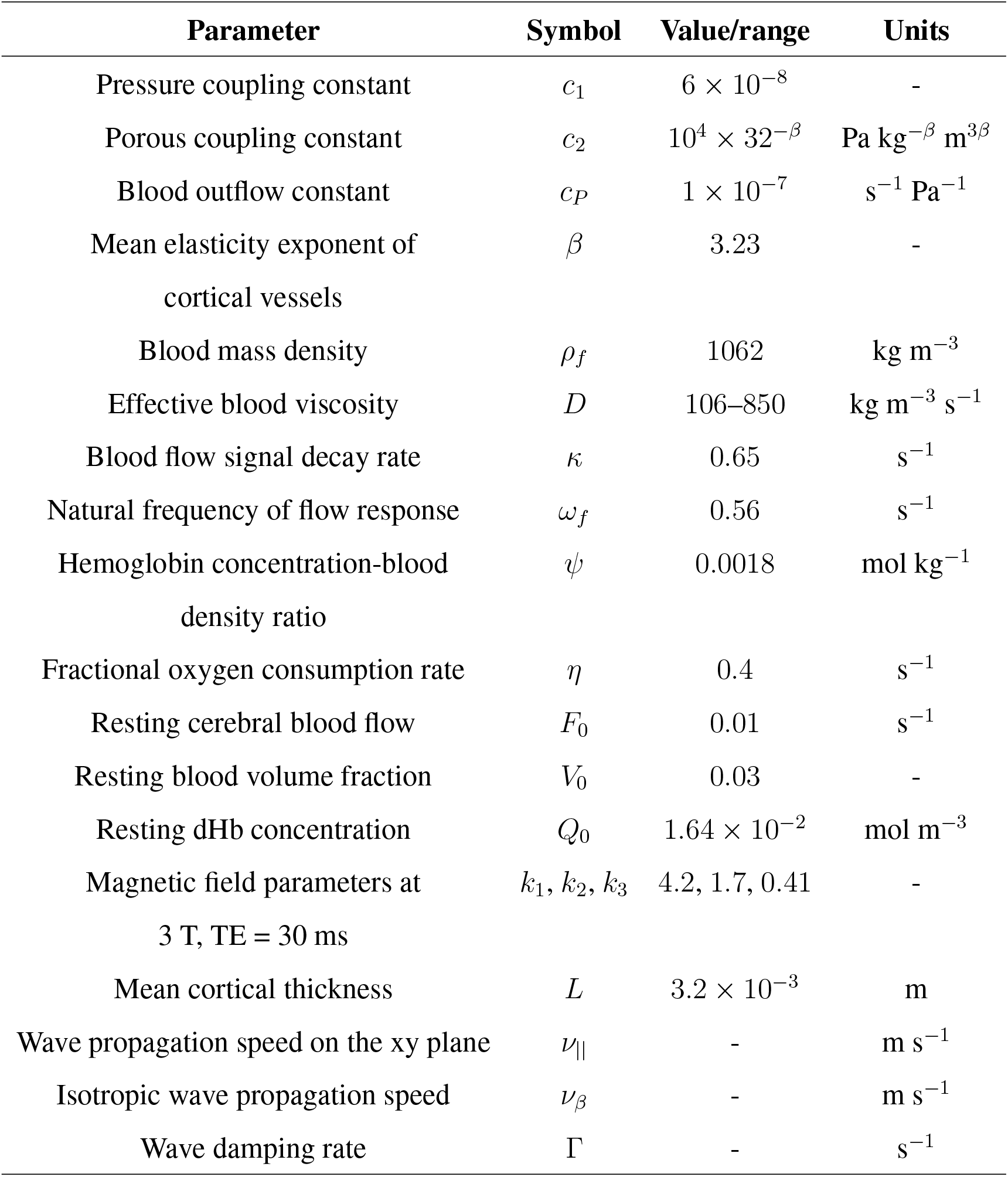
Model parameters. In each row, columns are ordered from left to right to detail the variable, its symbol or formula, its value or range, and its units, respectively.

### 2.2. Modeling Procedure

Because we want to compare the full 3D model directly to the results of Puckett et al. (2016) who used a 2D layered model, which is discussed in more detail below, we model the same neural drive as in their paper. The visual stimulus used in Puckett et al. (2016) was a thin, black and white flickering ring, which induces a strong response for all parts of the primary visual cortex (V1) that respond to a particular eccentricity in the visual field. Because the radial eccentricity of the stimulus was fixed, the retinotopically mapped responses induced approximately a straight line in V1. Hence, the response perpendicular to the line will be approximately constant along the line’s length due to the translational symmetry of the system (Aquino et al., 2014). This means that when simulating the stimulus, we can take advantage of this symmetry and ignore the direction parallel to the stimulated line, leaving only two spatial dimensions, i.e., *z* (for the direction corresponding to the depth) and *x* (for the remaining direction perpendicular to *z*). This significantly hastens the computations and allows for more accurate simulations by allowing more computational memory to be devoted to the other spatial dimensions. However, for more general stimuli, this approximation cannot be made and it is necessary to model all three spatial dimensions.

To try and reproduce the results of Puckett et al. (2016) accurately, while taking into account the full 3D nature of the system, we apply a 4 s visual stimulus corresponding to a single point in the *x*-dimension, with the blood inflows in the *z*-dimension a Gaus-sian centered at approximately *z* = 2.4 mm (the same as the inflow 0.8 mm below the surface discussed in Sec. 2.1) with a FWHM in *z* of approximately 1 mm, resulting in most blood inflows being confined to near the surface. A closed boundary is applied at *z* = 3.2 mm (because blood can’t flow through the cortical surface), and also at *z* = *-*0.8 mm, in the white matter. All inflows and outflows occur within the main body of the tissue.

Schematic diagrams of the 2D and 3D models are shown in Fig. 2. The figure shows the main point of contrast, in that the 2D model in Fig. 2a has all blood flows confined in each *z* layer, while the 3D model in Fig. 2b allows for blood flows to occur in any direction. Furthermore, due to the layered property of the 2D model, the relative sizes of inflows in different layers must be input as a model parameter. On the other hand, the 3D model naturally predicts the changes in BOLD in each layer relative to each other by taking into account how far that depth is from the main inflow depth.

**Figure 2:**
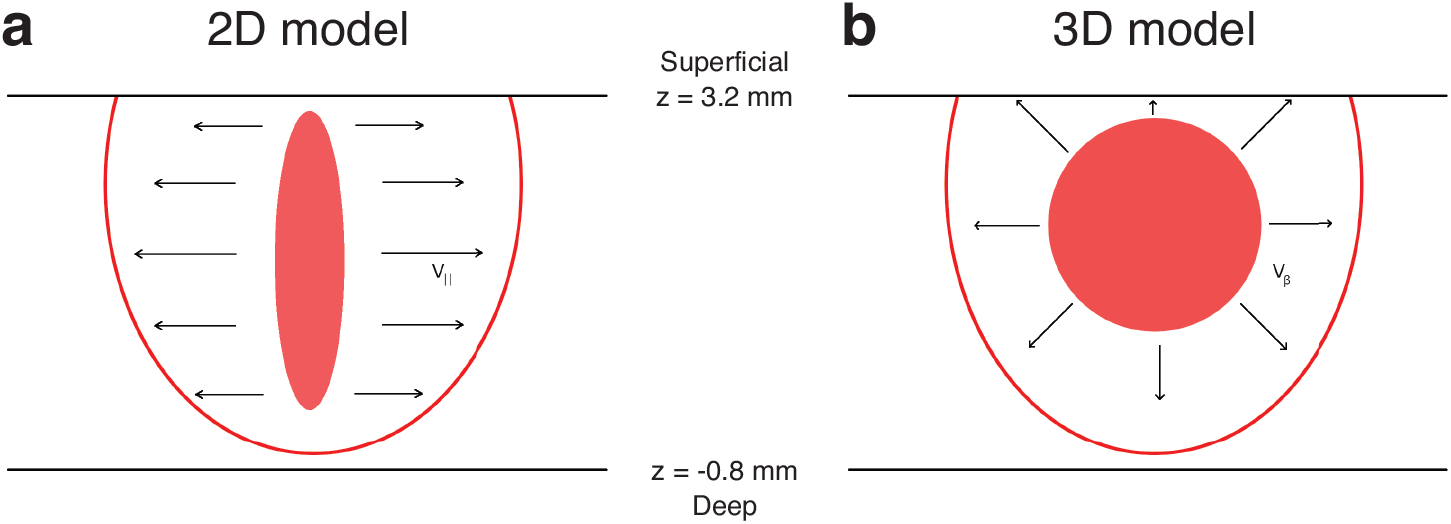
Schematic diagrams of 2D and 3D models of how blood inflows spread throughout the cortex. (a) 2D model used in Puckett et al. (2016), where inflows are confined to a fixed *z* and only propagate laterally. *v*_||_ is the velocity of propagation in a particular layer. (b) 3D model proposed in this work, where inflows are localized to a much smaller range of depths and blood is allowed to flow freely to different *z*’s. *v*_*β*_ is the general velocity of hemodynamic waves, which is the same in all directions, causing isotropic spreading.

The baseline value for the blood mass density Ξ changes with *z* due to the different numbers of blood vessels. Therefore, to include this in the model, we change the baseline Ξ according to the baseline percentage of blood vessels at each depth estimated by Duvernoy et al. (1981). Similarly, other properties of the cortical tissue (e.g., *β* that is related to the elasticity of the blood vessels) are likely to change with *z*, but these are more difficult to determine precisely due to them not being directly measurable. In fact, because some parameters are very difficult to determine noninvasively in general, fitting the modeled response to a measured BOLD signal could actually help narrow down the ranges of these properties in a subject via fMRI alone.

### 2.3. Experimental Setup

The main details of the experiment being compared to were discussed in Puckett et al. (2016), but are briefly outlined here. Six subjects had MRI scanning performed upon them with a Siemens MAGNETOM Trio 3 T MRI scanner. Several T1-weighted anatomical images were collected and then aligned through a rigid body transformation to allow for estimation of distances along the cortex. The functional data were obtained using a gradient-echo sequence with a matrix size of 240*×*240 and an FOV of 192 mm, resulting in an in-plane resolution of 0.8 mm *×* 0.8 mm.

Visual stimulation was done on a white screen viewed via a mirror. The stimulus itself was a ring stimulus with a checkerboard pattern at an eccentricity of 2 degrees, flickering every 250 ms. This was presented for 4 s, before being removed for 16.5 s. This cycle was repeated 8 times per run, giving a 3 min run, and 18–25 runs were collected per subject. Retinotopic maps were also obtained to ensure accurate mapping of the activity onto the cortex; this was done using a rotating bowtie stimulus, as in Schira et al. (2009), for 2 runs of 6 min each.

## 3. Results

In this section, we use the model discussed above to predict the BOLD responses to stimuli in the visual cortex. We then compare the modeled responses with the results of Puckett et al. (2016) and demonstrate its accuracy and advantages.

### 3.1. Model Predictions

Figure 3 shows the predicted BOLD responses of the 3D hemodynamic model for the same stimulus input as was used in the experiment by Puckett et al. (2016). Each frame is an *x*-*t* cross-section of the measured response at a different *z* (going from the surface at *z* = 3.2 mm to *z* = *-*0.8 mm in steps of 0.8 mm, with the gray-white matter boundary at *z* = 0 mm). The general pattern at each *z* is an initial increase in the induced response as the stimulus is applied, reaching a maximum after approximately 5 s, before decreasing rapidly and giving an undershoot in the BOLD signal that peaks at approximately 11 s in each frame (0 s is defined as the start of the stimulus). The main difference between frames are the magnitudes of the positive peak and negative trough in the response, which are largest for *z* = 3.2 mm and decrease as one moves away from that depth.

**Figure 3:**
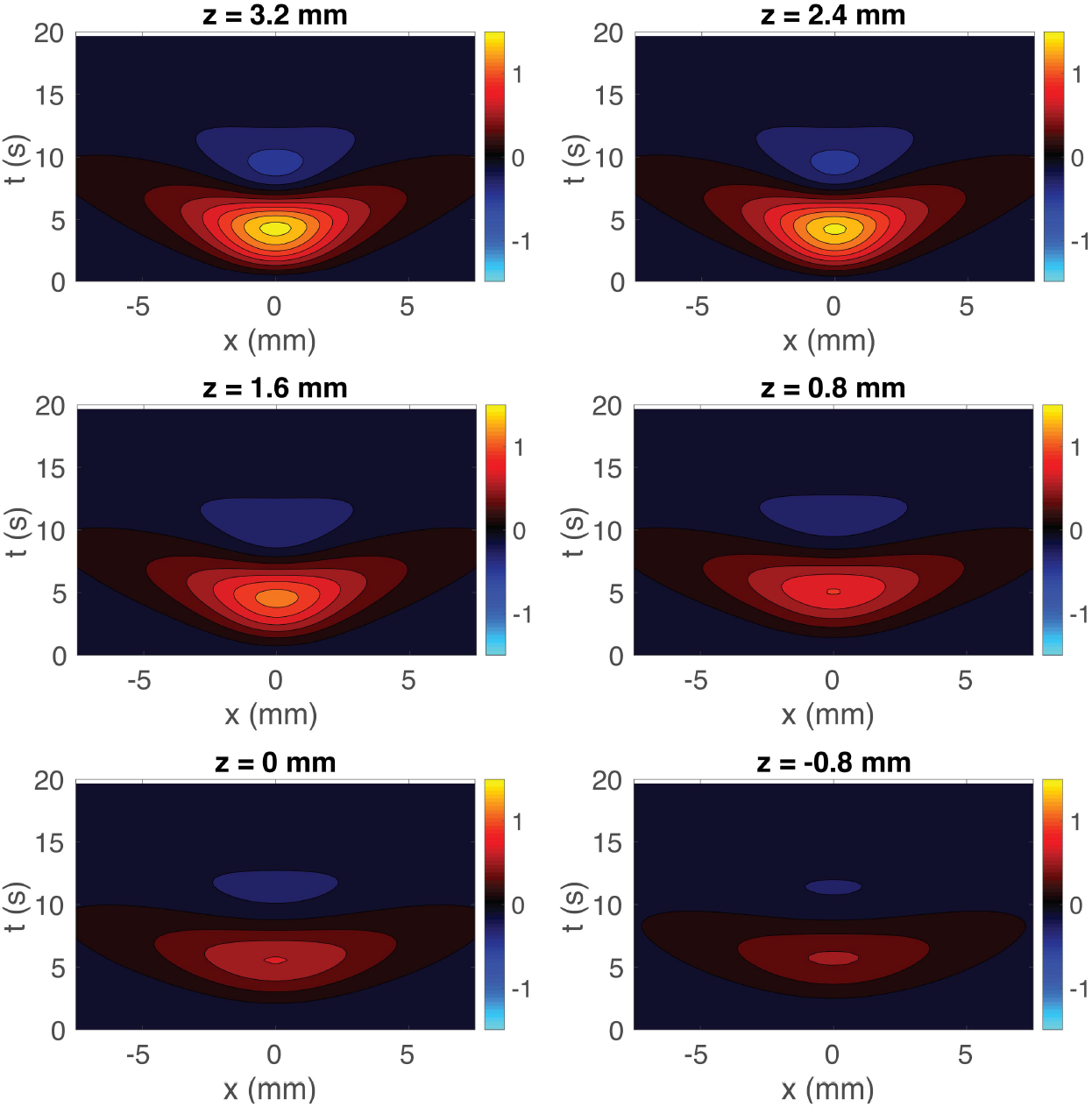
Cross-sections of the predicted BOLD response induced by a line stimulus using the 3D model. The responses are shown for *z* = 3.2, 2.4, 1.6, 0.8, 0, and *-*0.8 mm (as labeled) vs. transverse distance *x* and time *t*.

In addition to examining the different depths separately, we next show how the response appears through all the cortical depths as a function of time; the results are shown in Fig. 4. Each frame is an *x*-*z* cross-section of the measured response at a fixed time from 1 to 8 s.

**Figure 4:**
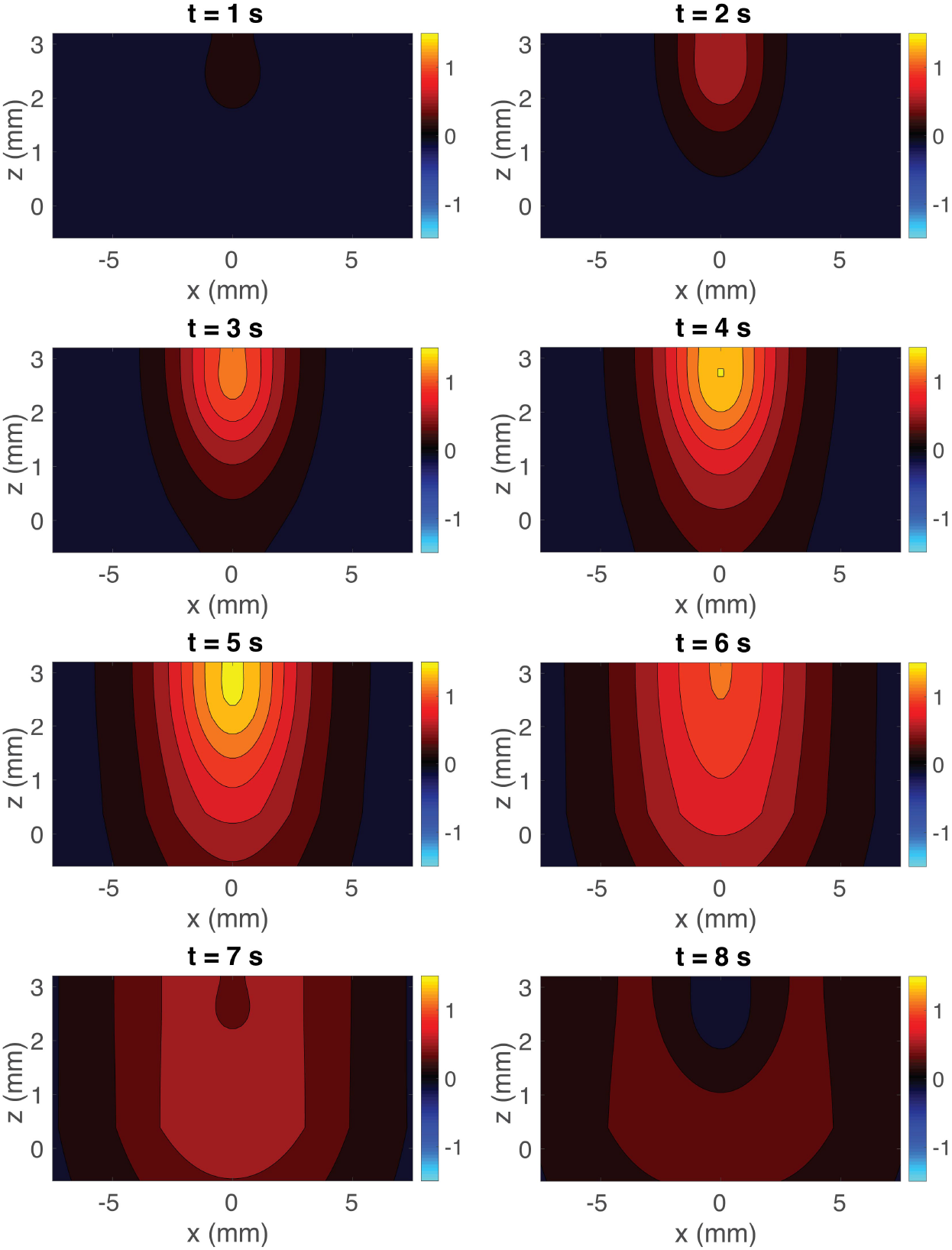
Cross-sections of the predicted BOLD response vs. the transverse distance *x* and the depth *z* at times from 1 to 8 s (as labeled) using the 3D model. The color scale is the same as in Fig. 3.

Figure 4 shows that the response starts off increasing mostly near the cortical surface (*z* = 3.2 mm), before spreading deeper and across the cortex. Eventually, when the stimulus is removed, the response decreases and then undershoots. This demonstrates how the changes in BOLD signal deeper in the cortex can be caused by blood flow changes closer to the surface, due to spreading hemodynamic waves.

### 3.2. Experimental Data

We now compare the simulated response with the experimentally measured response from Puckett et al. (2016) shown in Fig. 5. The experimental response are formatted similarly to Fig. 3, which shows the induced BOLD response averaged over all 6 subjects (Fig. 5 left) and for one particular subject (Fig. 5 right) due to a 4-s line stimulus. At all depths there is an initial increase in the BOLD response near to where the stimulus is applied, followed by spreading into nearby areas. Moreover, the response tends to decrease as *z* decreases going closer to the gray-white matter boundary. However, the experimental results have slight differences compared to Fig. 3, with regard to which *z* depth has the highest response and the exact time taken to reach a maximum response, but these are small and can be explained by small differences in physical parameters such as *β* between subjects [it is well within the variation between subjects seen in Puckett et al. (2016)].

**Figure 5:**
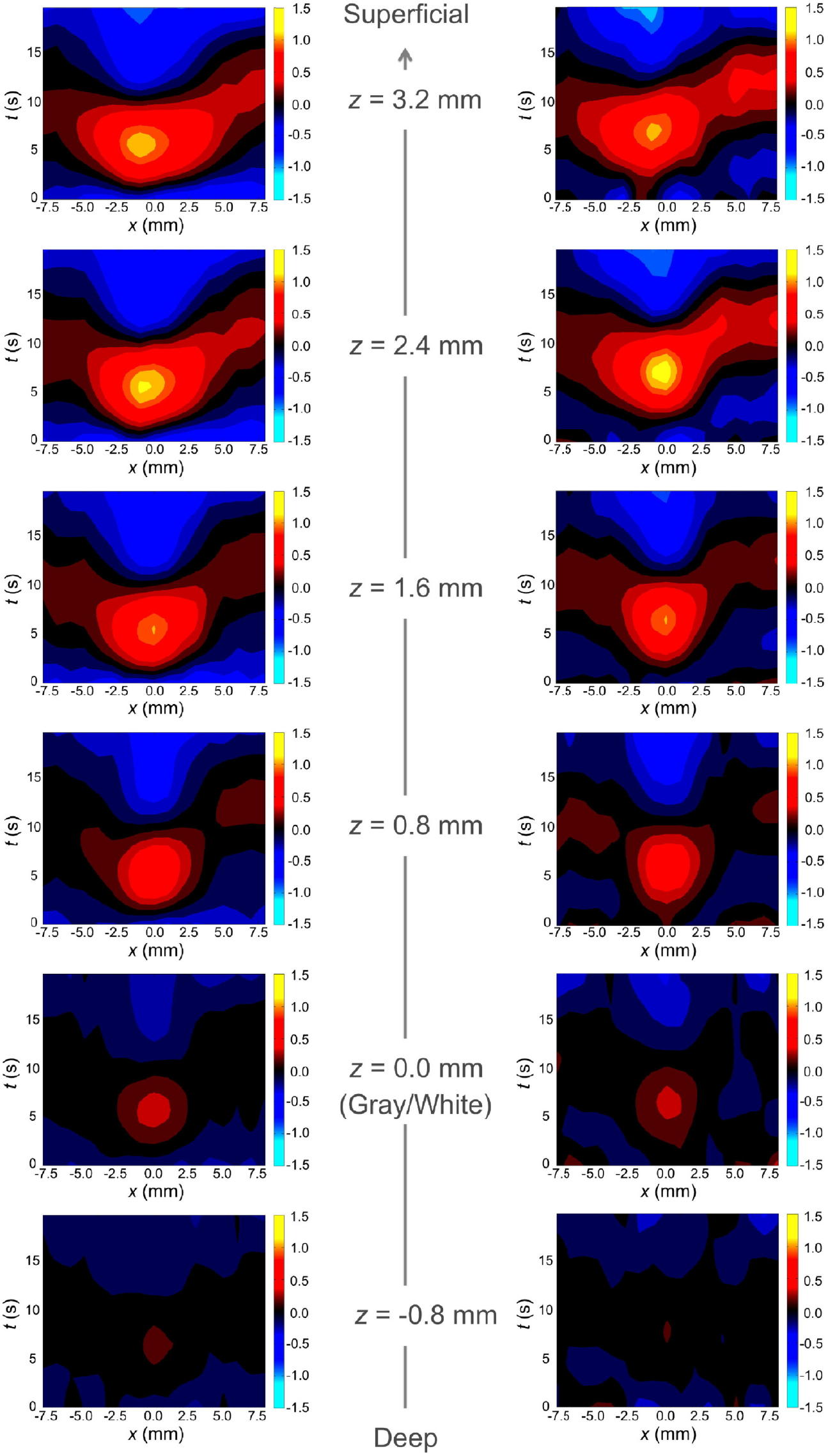
Group-averaged (left) and single-subject (right) experimental BOLD responses relative to the distance from the gray-white matter boundary at *z* = 0 mm. The color scale is the same as in Fig. 3. Figure reproduced from Puckett et al. (2016).

We then compare our results with those predicted using the 2D model, where each depth is considered as a separate 2D system [similar to the modeling results in Puckett et al. (2016)]. For the 2D model in Puckett et al. (2016), the spatial spreading, the peak amplitude, the hemodynamic velocity, and the damping rate were estimated for each depth from the experimental data in Fig. 5. These estimates are shown in Fig. 6, where a linear fit vs. depth is shown for each. The fits were used to make 2D model predictions at each depth, thereby yielding Fig. 7. This shows the predicted spatiotemporal response in the same format as the experimental response in Fig. 5, with the spatial spreading, amplitude of response, and hemodynamic velocity being adjusted at each depth to fit the data.

**Figure 6:**
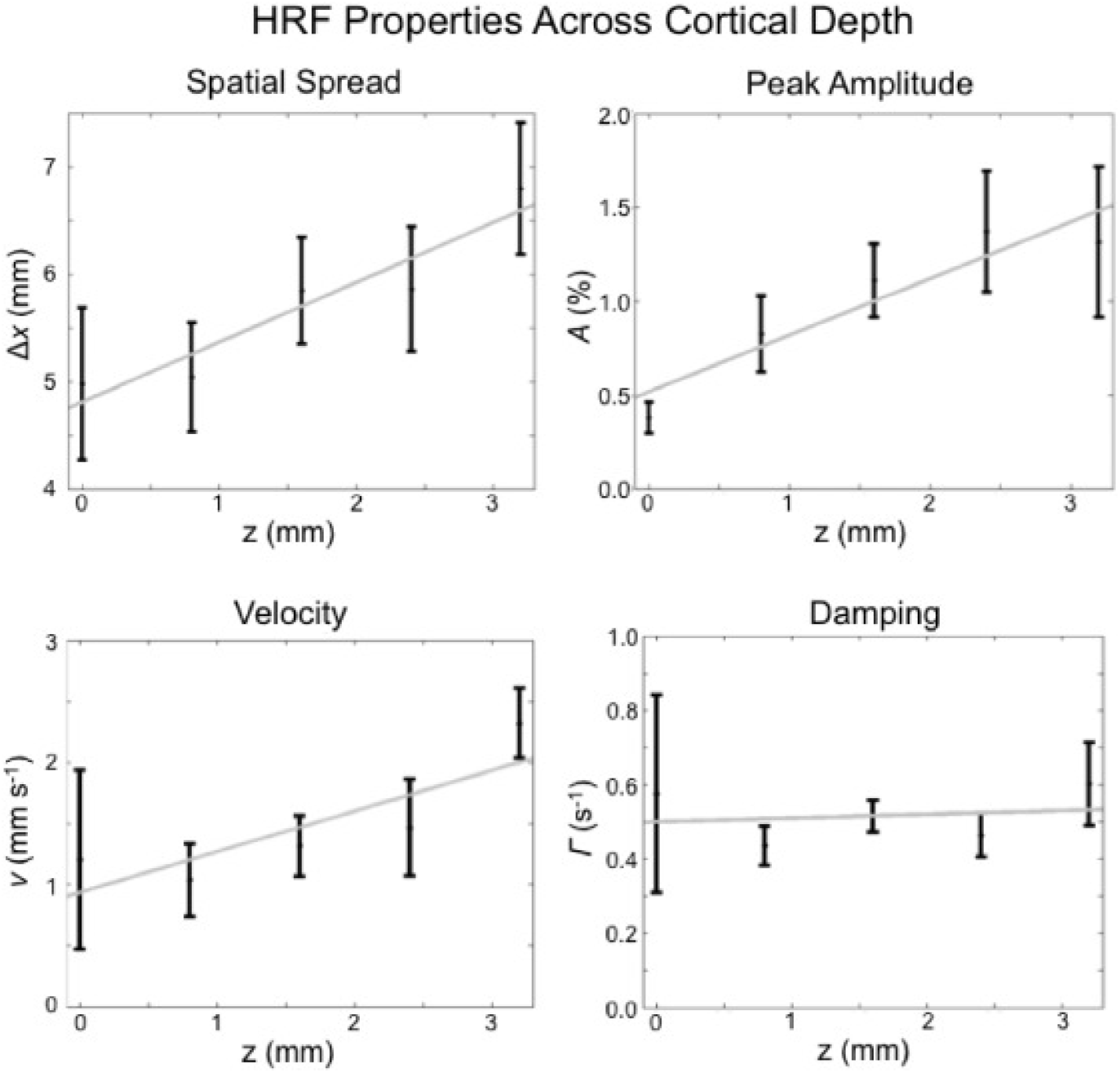
Measured values of some properties of the induced hemodynamic response versus the cortical depth. The properties considered are: spatial spread *δx*, peak amplitude *A*, velocity *v*, and damping Γ. Note that the velocity here corresponds to *v*_||_ in Fig. 2a. The dots represent mean values and the vertical lines represent standard errors. The gray lines represent linear fits. Figure reproduced from Puckett et al. (2016).

**Figure 7:**
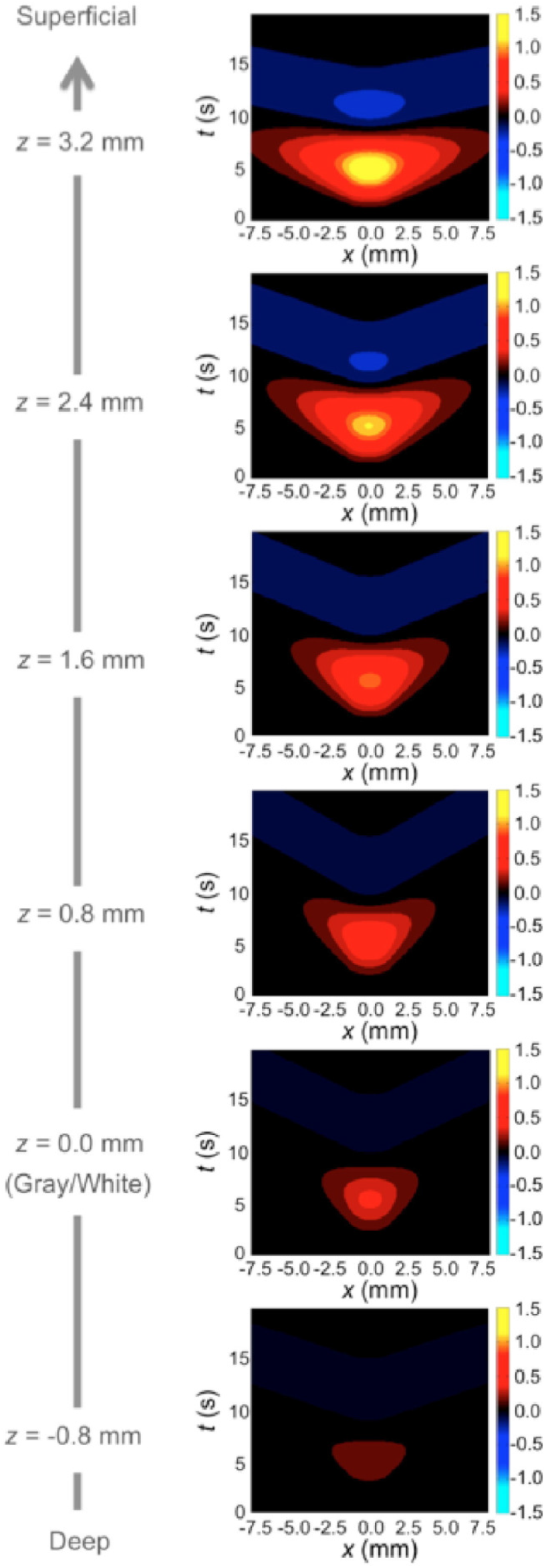
Predicted BOLD response using the 2D model. The maximum BOLD response in each *z* layer is adjusted to lie on a Gaussian with maximum at *z* = 3.2 mm.

The results in Fig. 7 reproduce those in Fig. 5 closely, with the same overall shape, time to peak, and decreasing amplitude as *z* decreases. However, many free parameters are involved in fitting the 2D layered model to the experimental data. Specifically, Fig. 7 involves linear fits of each of the 4 parameters shown in Fig. 6, giving 8 degrees of freedom, so it is unsurprising that a good fit was obtained this way.

### 3.3. Comparison of 3D model to 2D layered model

Figure 3 shows a close reproduction of the responses in Fig. 5 using our conceptually simpler and physically more realistic 3D model. However, to show that it is a significant improvement over the previously used 2D layered model, we need to compare the predictions more directly. The main prediction of the full 3D model is that blood flows isotropically out of a localized inflow area instead of inflows being located at all depths. Hence, we rescale Figs 4 and 7 so that they are in the same format, and likewise reformat the experimental data in Fig. 5 to show the time-varying cross-sectional *x*-*z* responses (see supplementary video at https://ars.els-cdn.com/content/image/1-s2.0-S1053811916302543-mmc1.mp4) so that it is directly comparable.

The results of the above rescalings are shown in Fig. 8, where Fig. 8a shows the predictions of the 3D model, Fig. 8b shows the predictions of the 2D layered model, and Fig. 8c shows the experimental data. Each column shows a gradual increase in BOLD signal over several seconds before peaking at about 5–6 s, and then a decrease before undershooting. In addition, all show the spreading of the response out from *x* = 0, with the response decreasing with distance. However, there are also significant differences, primarily between Fig. 8b and the other two. In the 3D model in Fig. 8a, the response spreads isotropically from the initial main inflow location because there is no preferred direction for the blood flow. This is very similar to what occurs in the experiment in Fig. 8c, which is also consistent with isotropic spreading. However, in the 2D model in Fig. 8b, the extent to which the response spreads in the *z*- and *x*-directions are not the same because the spatial spreading in each direction is a free parameter. Given how the 2D layered model incorporates the variation in parameters vs. *z*, there is no prior reason for the wavefronts to appear to be approximately spherical. This behavior is easy to explain with the 3D model due to the spreading having no preferred direction, thus explaining more properties of the system while having fewer assumptions and parameters. The behavior of the response for deep *z* is also of interest for future studies because the 3D model predicts that there should be a detectable wave induced there 1–2 s after a short stimulus due to a time delay for hemodynamic flows to reach small *z*, which is not predicted by the 2D layered model where changes in the source are simultaneous at all depths.

**Figure 8:**
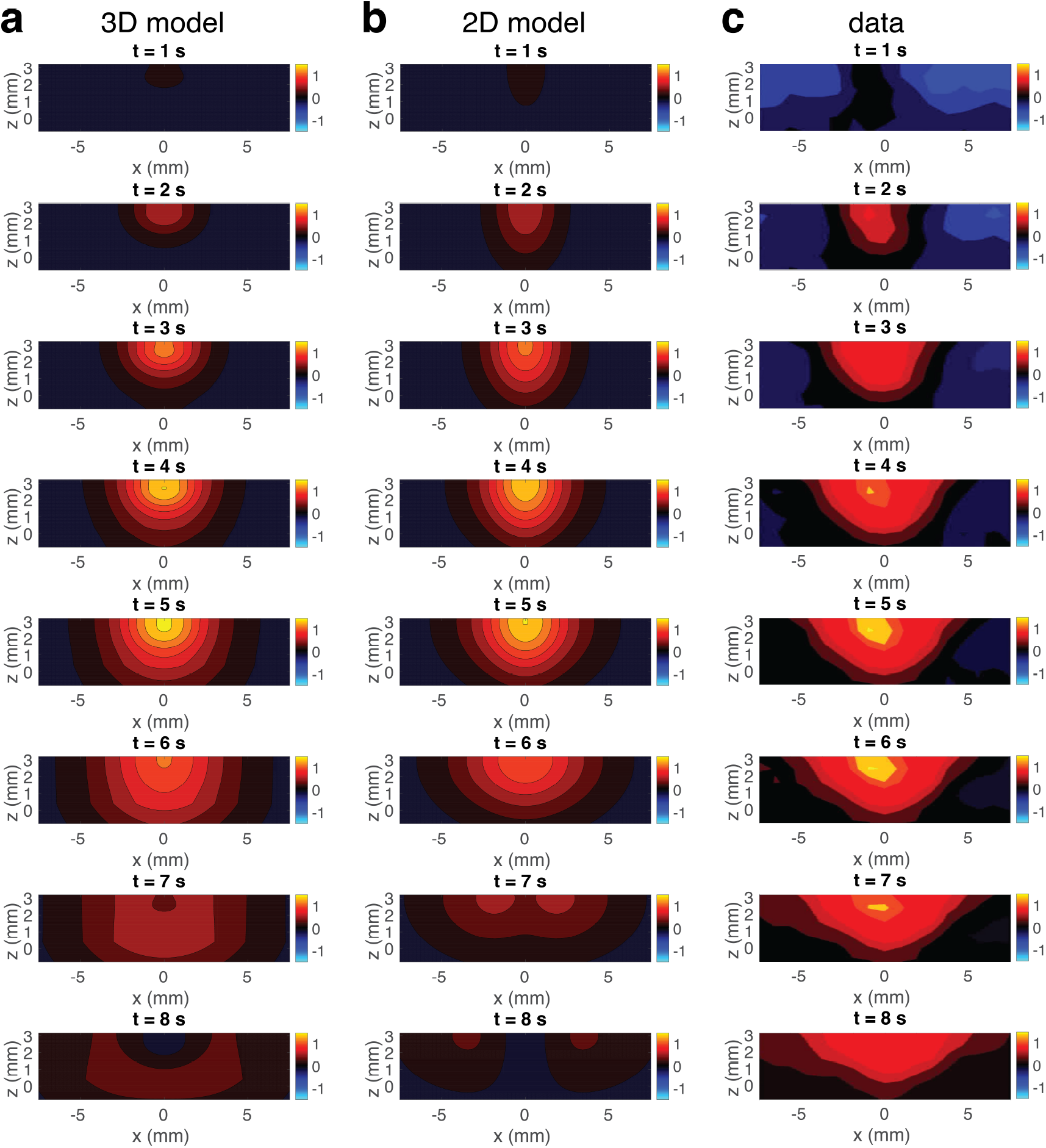
Predicted and experimental BOLD responses. (a) Predicted responses using the 3D model. The panels are the same as in Fig. 4 but with the axes adjusted to remove spatial distortion. (b) Predicted responses using the 2D layered model. The responses are produced by setting model parameters that match those used in Fig. 4 of Puckett et al. (2016). (c) Experimental responses from Puckett et al. (2016). For a, b, and c, the rows are arranged from top to bottom to show the responses from *t* = 1 to 8 s, respectively. Moreover, the color scale for each panel is the same as in Fig. 3.

In addition to explaining the isotropic spreading of the BOLD response, we also examine the parameters used in the 2D layered model shown in Fig. 6 to determine whether their variation with *z* can also be explained by the 3D model. In the 3D model, the physical parameters are independent of *z* (apart from a change in *β* introduced near the gray-white matter boundary). However, Fig. 6 shows that there are discernible changes in several of the hemodynamic properties of the 2D layered BOLD signal as *z* varies, except for the damping rate Γ. The explanation of these changes is that the variations of these inferred parameters of the 2D layered model are caused by fitting them to mimic the actual isotropic 3D blood flows. At depths far from the inflow depth, the separation between the inflows and the point being considered is significantly larger than the distance to the *x* = 0 location at that depth. By Pythagoras’ theorem, we have

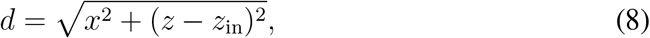

where *d* is the physical separation of the point at (*x, z*) from the primary inflow source and *z*_in_ is the depth of inflows. This means that for *z ≈* 0, which is near the gray-white matter boundary, *d* will be significantly larger than *x*, and so this means the apparent values of most parameters in Fig. 6 (i.e., the 2D layered parameters that most closely mimic the 3D effects) will change as a result. As the method of determining the spatial spreading in Puckett et al. (2016) is to assume a Gaussian profile, the spatial spreading will be adjusted by a factor of *x/d* as the distance measured in the 2D system is *x*, while the separation in 3D space is *d*. Given the isotropic spreading of the response, the peak amplitude of the response in each layer of the 2D model, *A*, should follow the same profile as the BOLD response in the *x*-direction. Finally, the hemodynamic velocity *ν* will appear smaller by a factor of *x/d* due to the delay induced by the response propagating diagonally through *z* as well. This is complicated by the effect shown in Fig. 9, where due to an initial delay in the wavefronts reaching deep *z*, the initial apparent horizontal velocity of the wave can be significantly higher than *v*_*β*_, with it then appearing to slow down as it moves to larger *x*. Each wavefront has a regular spacing and is moving at a constant speed outward from the source, but the apparent motion along the *x*-direction can be significantly higher and vary due to how the spherical wavefronts intersect a horizontal plane. This is seen clearly in Fig. 9, as the distance between the intersection points of the circular waves and the planes in the *x*-direction (shown for the highest layer with a blue line) are significantly larger than the separation of the waves along a line perpendicular to all the wavefronts (shown by a red line). However, for Puckett et al. (2016), the fitting used to determine the velocity was done from the peak of the BOLD response in each layer, which should mostly mitigate this effect (it will be strongest at the leading edge of the wavefronts). Most importantly, we conclude that both Δ*x* and *ν* will vary with *x* in the 2D model.

**Figure 9:**
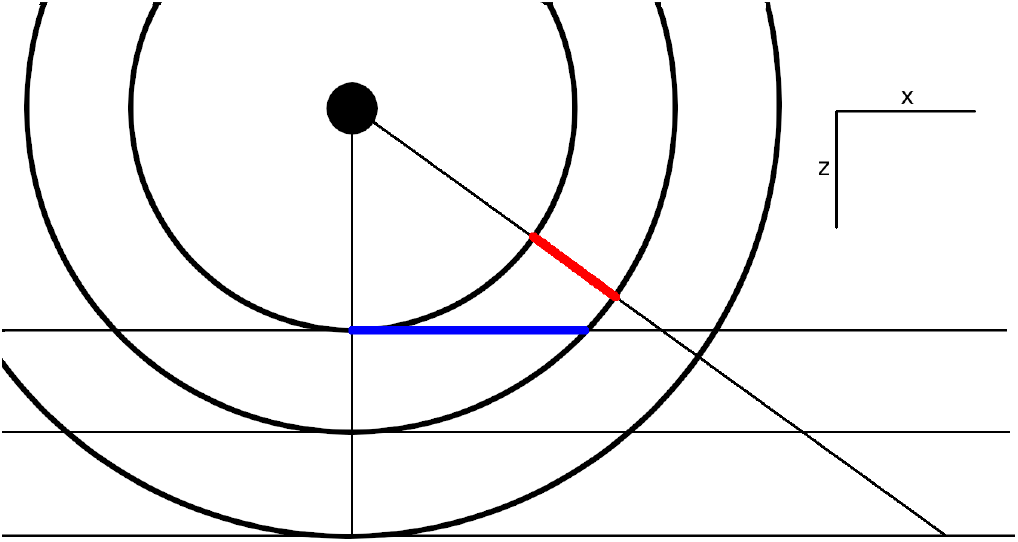
Illustration of how the apparent velocity of a wave along a plane can vary if not measured along the direction of travel. The blue line shows the apparent separation of two wavefronts as observed along the highest *z* layer, while the red line is the actual distance the wave has travelled. The *x*- and *z*-directions are labelled on the right.

In Puckett et al. (2016), the spatial spreading Δ*x* (the FWHM of the response, fitted as a Gaussian) was calculated by determining the FWHM of the response when the BOLD response peaked, while *ν* was calculated using the method from Aquino et al. (2012) by fitting lines from *x* = 0 at the time when the response peaks to outgoing wavefronts at least 2 mm from the localized response. Hence, using this information, and estimates of Δ*x, A*, and *ν* at *z* = *z*_in_, we use the full 3D model to infer the 2D parameters that will be needed to mimic the 3D effects, which gives Fig. 10. In this figure, we can see the data for each of the 2D layered values of Δ*x, A*, and *v*, along with the curve corresponding to the values of each parameter predicted using the isotropic 3D model. We see that the predictions for Δ*x* and *A* are very close to the observed values, while the predictions for *ν* fit most of the depths well, except perhaps at the *z* = 3.2 mm point (which could be due to the hard boundary at the cortical surface requiring the blood flow to accelerate as it is constricted). These accurate predictions demonstrate that most of the variations in these 2D layered parameters can be explained as resulting from evoked 3D blood inflows being localized to a specific depth near the surface of the cortex, rather than strong increases in evoked multilayer inflows along with layer-dependent hemodynamic properties as well.

**Figure 10:**
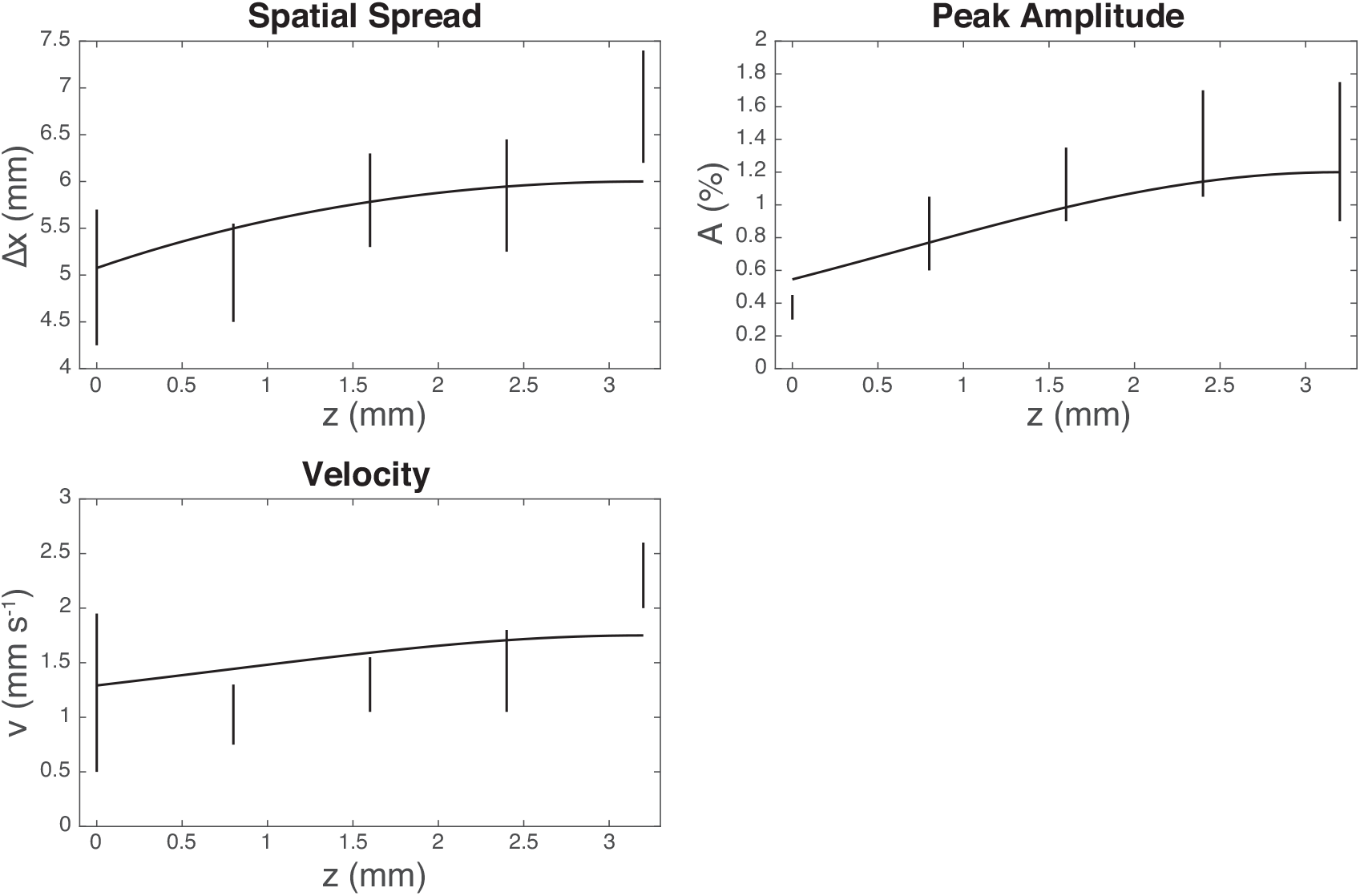
Same as Fig. 6 but now includes predicted curves, assuming that the the variation is due to effects induced by 3D blood inflows.

## 4. Summary and Discussion

A nonlinear 3D spatiotemporal model of hemodynamics for predicting induced BOLD response was introduced, thereby relaxing assumptions made in earlier works such as the previous 2D layered model used in Puckett et al. (2016). The new model was solved numerically and the results were compared to data from the experiments of Puckett et al. (2016) and with parameter estimates obtained from those experiments under a 2D layered approximation. The main results are:

i. The decrease in BOLD amplitude and spatial spread at depths farther from the cortical surface can be explained by the main blood inflows due to the neural drive localized to *z* near the cortical surface, with hemodynamic changes propagating isotropically, including down through the cortex, as predicted by the 3D hemodynamic model.
ii. The main features of the induced BOLD response to a line stimulus in V1 can be accurately reproduced via the 3D physiologically based model. Using the full 3D model increases predictive power, while simultaneously decreasing the number of parameters and assumptions required compared to the 2D layered model. For example, there is no more need to define depth-specific parameter values for spatial spreading, peak amplitude, and hemodynamic velocity.
iii. The observed apparent changes in properties of the BOLD response by the 2D layered model such as the transverse spatial spread Δ*x* and apparent hemodynamic velocity *ν* can be explained, within standard error, by the effects of trying to treat properties of a 3D system by examining 2D slices of that system. The transverse components of the spreading and velocity appear to be lower farther from the surface in the 2D model due to the larger separation points at that depth from the inflow source, unless this effect is specifically corrected for. No such correction is necessary in the 3D model and the experimental results are consistent with isotropic spreading.

Future experiments could explore the effects of the isotropic spreading of the BOLD response throughout the cortex by designing stimuli appropriately and imaging the entire thickness of the cortex at high resolution, via advanced laminar fMRI protocols (Norris and Polimeni, 2019). In particular, if a short stimulus is applied and then removed, a 2D layered model would predict that the response deep in the cortex would peak simultaneously with the response near the surface. However, the 3D model predicts that a sufficiently short stimulus will induce a wave traveling isotropically within the cortical sheet from the source depth, and so should lead to a measurable time delay on the peak BOLD response for deep layers. Observing this directly would conclusively show that the changes in BOLD signal with depth are primarily due to vertical transmission of the hemodynamic waves. However, care would need to be taken to avoid any possible nonlinearities in the response to short, strong stimuli required to observe such an effect (Birn and Bandettini, 2005), although these would not be likely to affect its isotropy.

Finally, MATLAB codes to simulate the model in this study are available at https://github.com/BrainDynamicsUSYD/hemodynamics-layers.

## 5. Acknowledgments

This work was supported by the Australian Research Council Center of Excellence for Integrative Brain Function (Grant CE140100007) and the Australian Research Council Laureate Fellowship (Grant FL140100025).

